# Preferred visual experiences provide intuitive descriptions of the functional properties of the cortical navigation network

**DOI:** 10.64898/2026.09.02.749001

**Authors:** Tianjiao Zhang, Cheol Jun Cho, Jack Gallant

## Abstract

Complex, high-dimensional brain representations are often difficult to understand intuitively. Recent advances in generative neural networks have made it possible to predict the optimal stimulus that maximizes the activity in a specific brain region. Because these stimuli are in image space, they provide an intuitive understanding of the functional properties of that region. Previous work on optimized stimuli has focused on passive perceptual tasks. Yet real-world perception operates in a closed loop with cognition and action, and so passive stimuli are unlikely to be optimal for many brain regions. In a previous study, we used fMRI to record BOLD activity from participants actively navigating through a virtual world. We identified a network of 11 cortical regions that support active navigation and are tuned for complex feature combinations. Here, we used a variational autoencoder to generate video clips predicted to correspond to maximal activity in each region. These preferred visual experiences capture dynamic scenes that incorporate both world events and participant actions. These preferred visual experiences provide intuitive descriptions of the functional properties of each region, and suggest that these regions are more engaged during dynamic than static scenes. These patterns also generate novel, data-driven hypotheses for future study.

## INTRODUCTION

Computational models of brain activity describe the tuning of the brain to stimulus and task features (Naselaris et al., 2011), yielding detailed descriptions of brain function. However, it is often difficult to understand brain function because tuning is often high-dimensional (Fusi et al., 2016; Rigotti et al., 2013). Additionally, beyond the primary sensory cortices, the brain is often selective for features in abstract spaces that are difficult to describe in intuitive terms (Bao et al., 2020; Huth et al., 2016). This issue is exacerbated in data recorded during complex naturalistic tasks, in which brain regions can be simultaneously selective for many task parameters (International Brain Laboratory et al., 2025; Zhang et al., 2025).

One method to overcome this interpretation challenge is to reconstruct the stimulus that is predicted to elicit the most brain activity. Early work in this direction combined voxelwise encoding models with a prior sampled from large libraries of natural images or video to reconstruct what a participant was viewing (Naselaris et al., 2009; Nishimoto et al., 2011). Recent advances in generative deep neural networks (Goodfellow et al., 2020; Gui et al., 2023) described below have replaced the explicitly sampled prior with a learned model of the stimulus space. Because a generative neural network can map any vector in an embedding space back into the original input space, an encoding model weight vector fit in that embedding space can be passed directly to the neural network to reconstruct the stimulus that maximizes the activity of a brain region. No search over a stimulus library is required, and the resulting stimulus provides an intuitive description of brain function as captured by the model.

Generative approaches to understanding the brain have been applied to both neurophysiology (Bashivan et al., 2019; Higgins et al., 2021; Ponce et al., 2019) and neuroimaging (Dado et al., 2022; Gaziv et al., 2022; Horikawa & Kamitani, 2017; Lin et al., 2022; Luo et al., 2023; Ozcelik & VanRullen, 2023; Ren et al., 2021; Shen et al., 2019; VanRullen & Reddy, 2019) data. Some studies have found good agreement between the reconstructed images and the reported functional properties of brain regions (Kamitani et al., 2025). Others have shown that these generated stimuli can be optimized iteratively to increase evoked brain activity, thus refining the stimuli across multiple rounds of presentation (Bashivan et al., 2019; Ponce et al., 2019).

However, prior generative modeling efforts have focused on stimuli in passive perceptual tasks. In contrast, real-world experiences are dynamic and closed-loop, shaped by both a continuously changing world and by our own actions. Previous work has demonstrated that tuning in the brain is strongly influenced by attention or task contexts (Çukur et al., 2013; Shahdloo et al., 2022; Zhang & Gallant, 2026). Thus, stimuli that are merely optimized for passive perceptual tasks may not necessarily reflect brain function during closed-loop naturalistic tasks. Furthermore, although one study showed that the latent features learned by an autoencoder resembles the selectivity of inferior temporal neurons (Higgins et al., 2021), few of these studies have used the generated stimuli to describe or interpret the functional properties of the brain.

To address these gaps, we use a variational autoencoder (VAE) (Kingma & Welling, 2019) to reconstruct dynamic video clips that describe the functional tuning properties of the human brain during naturalistic navigation. In previous work, we used fMRI to record brain activity from participants performing a taxi-driver task in a virtual city (Zhang et al., 2025). High-dimensional encoding models identified 11 cortical regions that support active navigation, and showed that these regions are tuned to complex combinations of visual-, navigation-, and motor-related features. To more intuitively understand the functional properties of these navigation-related regions, we used the VAE to reconstruct video clips that the fit encoding models predict will correspond to maximal activity in a brain region. Because the term “stimulus” implies that the visual scene is purely a driver of brain activity and is not influenced by participant actions, here we instead refer to the reconstructions as “preferred visual experiences” that capture the visual aspect of the closed loop. We then used these preferred visual experiences alongside the encoding model results to interpret the functions of the 11 regions in the cortical navigation network.

## RESULTS

In an earlier study we used voxelwise encoding models to identify a network of 11 functional regions in the cerebral cortex that support active navigation (Zhang et al., 2025). The functional properties of these regions are high-dimensional and difficult to interpret intuitively (Fig. 1A). To gain a better understanding of their functional properties, we used a VAE to reconstruct video clips that correspond to their preferred visual experiences. To do so, screen recordings from the navigation experiment were downsampled to 120×90 pixels at 3.75 frames per second, and a VAE with a 512-dimensional latent space was fit to these 2-second video clips (Fig. 1B). Voxelwise encoding models for activations in this latent space were fit to brain activity and explain activity broadly across the cortex (Fig. 1C, for performance in individual regions in the navigation network, see Supplementary Fig. 1). To reconstruct the preferred visual experience that is predicted to correspond to maximum activity in each region, the average voxelwise encoding model weight vector for that region was then projected into video space by the VAE decoder (Fig. 1D). Complementing these preferred visual experiences, we also projected the negated weight vector into video space, reconstructing the scenes that are predicted to correspond to the least activity in each region. We refer to these reconstructions as the “anti-preferred visual experience” for each region. Inspection shows that the VAE reconstructs visual experiences that contain interpretable structure, including building-like textures, prompts, road lines, other vehicles, and, notably, intelligible speedometer values that enable direct inference of egomotion through the visual scene (Fig. 1D).

**Figure 1.**
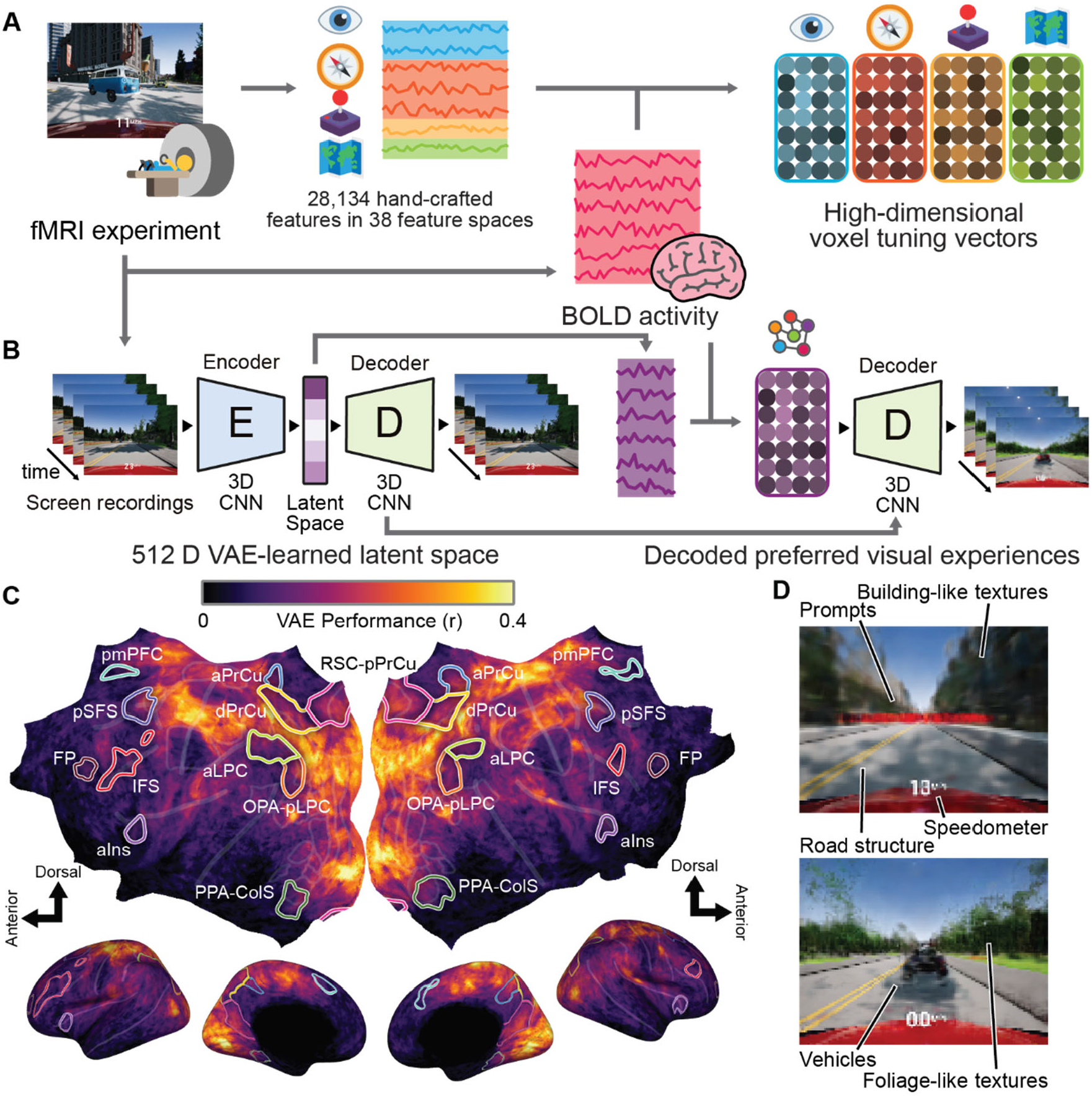
Variational autoencoder models support intuitive interpretation of brain activity. (A) Voxelwise encoding models provide a detailed description of brain function by describing its tuning to high-dimensional features, but the resulting complex tuning properties can be difficult to interpret intuitively. In naturalistic experiments, perception operates in a closed loop with cognition and action, so brain activity depends not only on perceptual input but also on internally-generated plans and the participant’s actions. (B) Variational autoencoders (VAEs) offer a data-driven solution to this interpretability problem. A VAE learns an embedding space for the stimulus, and its decoder projects any vector in that space back into stimulus space. Encoding model weights fit with the VAE embedding space can be given to the decoder to reconstruct the visual experience that is maximally correlated with activity in a voxel or brain region. These reconstructions, which we term “preferred visual experiences”, can be examined alongside encoding models to provide complementary, intuitive descriptions of brain selectivity. (C) We used this approach to interpret brain activity recorded during a naturalistic navigation experiment. Encoding models fit with the VAE latent space explain BOLD activity broadly across the cerebral cortex, indicating that the VAE latent space is well-aligned with brain representations. (D) The preferred visual experiences contain recognizable components, such as environmental structures, prompts, vehicles, and speedometer values. While the participants’ plans and actions that shape the scene are not reconstructed directly, the VAE reconstructs their visual correlates and enables detailed interpretation of functional tuning.

### Preferred visual experiences are plausible for regions with known functional properties

To confirm that the preferred visual experience reconstructions are meaningful and plausible, we first examined two regions whose functional properties have been well established by prior research: primary visual cortex (V1) and the primary hand motor area (M1H). The preferred visual experience for V1 consists of dense edges and shapes distributed across the screen, while the anti-preferred visual experience is dominated by open skies and empty roads (Fig. 2A). V1 is therefore most active when the participant drives through visually cluttered parts of the city, and least active in open areas that contain little visual structure. This pattern is consistent with the selectivity of V1 for low-level visual structure (Field, 1987).

**Figure 2.**
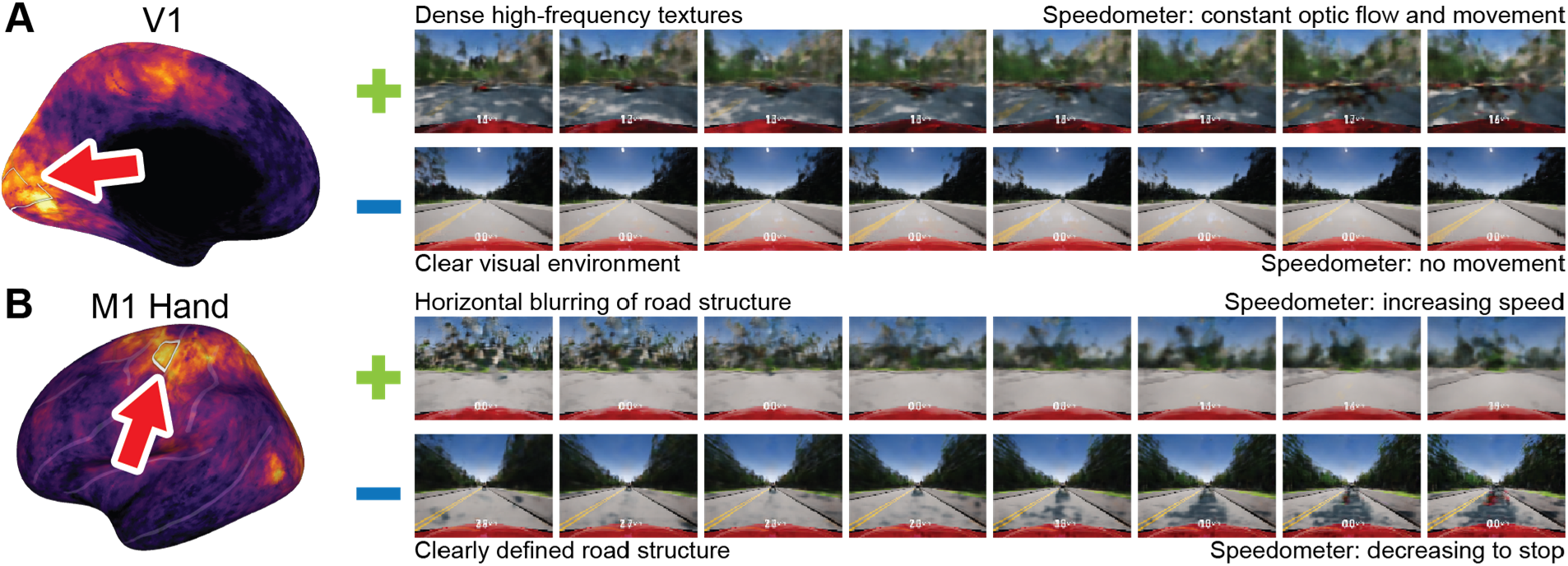
VAE encoding models recover plausible preferred visual experiences consistent with the properties of established functional regions. Encoding model weights for the VAE latent space are passed through the decoder to recover preferred and anti-preferred visual experiences for each brain region. While the preferred visual experience is correlated with the maximal activity of a region, the anti-preferred visual experience is correlated with the least activity in a region, and provides a contrast for interpretation. To confirm that the reconstructed preferred visual experiences are meaningful and plausible, we examined two regions whose functional properties have been well-established: primary visual cortex (V1) and the primary hand motor area (M1H). (A) The preferred visual experience for V1 consists of dense edges and shapes distributed across the screen, while the anti-preferred visual experience is dominated by open skies and empty roads. This pattern is consistent with the selectivity of V1 for low-level visual structure. (B) The preferred visual experience for M1H corresponds to turns that blur the street structure, while the anti-preferred visual experience depicts the vehicle coming to a stop. Because hand motor activity is used to steer the vehicle in this experiment, turning induces horizontal blurring of the visual stimulus. Together, these results confirm that VAE encoding models recover interpretable and plausible preferred visual experiences.

The preferred visual experience for M1H consists of horizontally blurred street structure, while the anti-preferred visual experience depicts the vehicle coming to a stop along a straight road section (Fig. 2B). Because hand motor activity steers the vehicle in this experiment, M1H is most active when the participant turns the wheel, which induces horizontal blurring of the visual input, and least active when the vehicle stops and no turning actions are possible. This pattern is consistent with the established role of M1H in hand movement (Penfield & Boldrey, 1937; Yousry et al., 1997). Both preferred visual experiences agree with the established properties of V1 and M1H, indicating that these VAE-derived preferred visual experiences are both interpretable and plausible.

### Preferred visual experiences for anterior visual navigation regions

Having shown that the VAE reconstructs plausible preferred visual experiences for well-characterized functional regions, we applied this analysis to the 11 cortical navigation regions identified in our earlier study (Zhang et al., 2025). We first examined three regions in the anterior visual cortex, the retrosplenial cortex-posterior precuneus region (RSC-pPrCu), the occipital place area-posterior lateral parietal cortex region (OPA-pLPC), and the parahippocampal place area-collateral sulcus region (PPA-ColS).

The preferred visual experience for the retrosplenial cortex-posterior precuneus region (RSC-pPrCu) includes the “go to” prompt, complex building-like structures, increasing speed, and blurring of the street structure (Fig. 3A). The anti-preferred visual experience depicts stationary periods in more open areas. RSC-pPrCu is thus most active at the beginning of trials, during rapid movement, and during heading changes in the city, and least active during stationary periods in open areas. This pattern builds upon our encoding models, which suggest that RSC-pPrCu transforms visual inputs into navigation representations. The preferred visual experience implicates RSC-pPrCu in scene representation, location updates, and heading transformations, and the presence of the “go to” prompt further suggests a role in route planning (Epstein & Vass, 2014) and orienting towards the goal (Vedder et al., 2017).

**Figure 3.**
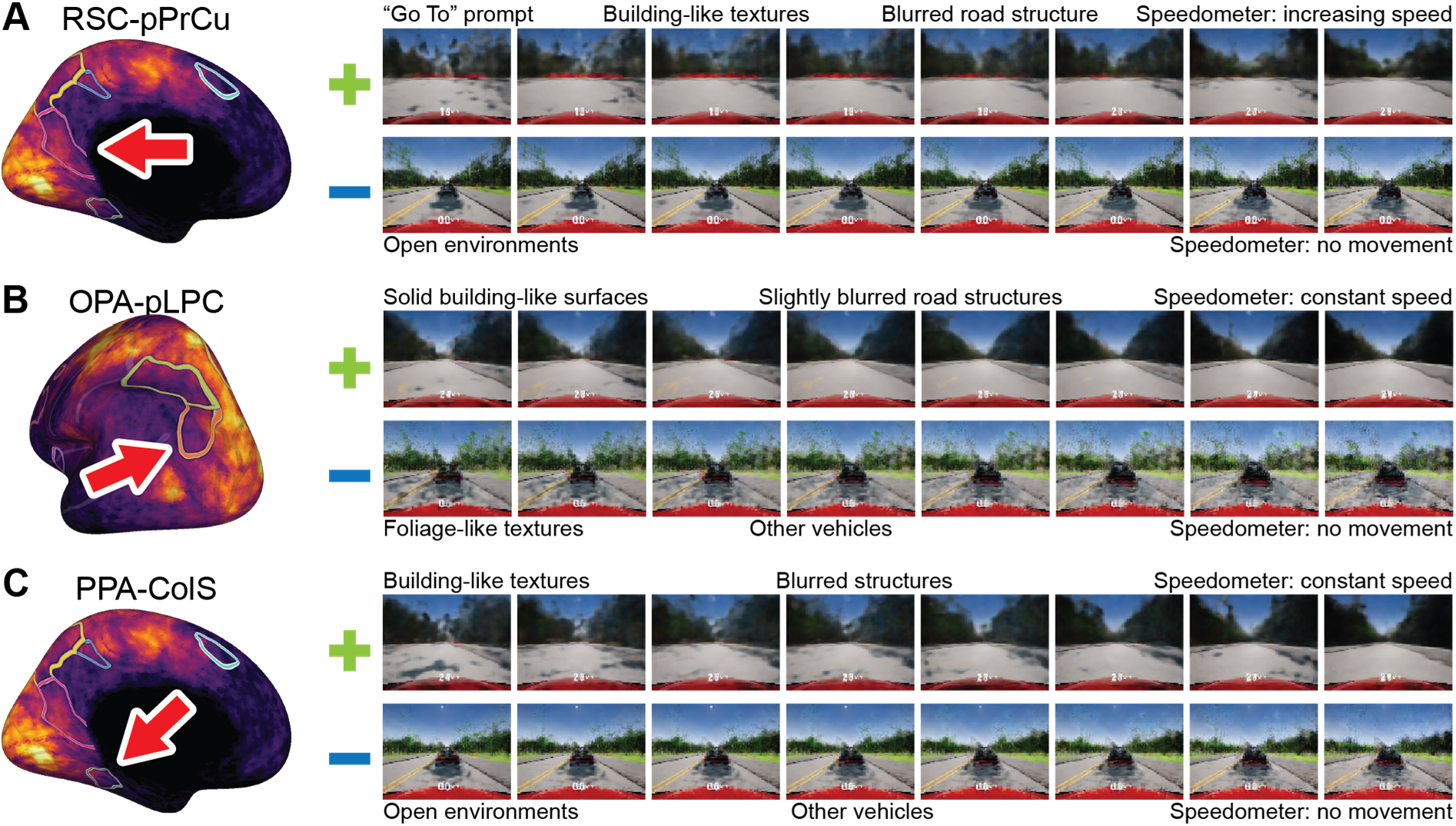
Preferred and anti-preferred visual experiences for anterior visual navigation regions. Our prior study revealed three regions in the anterior visual cortex, and suggested that these regions bridge perceptual inputs with the rest of the navigation network. (A) The preferred visual experience for RSC-pPrCu includes the “go to” prompt, complex building-like structures, increasing speed, and blurring of the street structure, indicating that the RSC-pPrCu is most active at the beginning of trials as the participant accelerates and turns through a dense urban environment. The anti-preferred visual experience depicts stationary periods in more open areas. This pattern suggests that the RSC-pPrCu may be engaged in transforming visual inputs to navigational representations and in route planning and orienting toward the goal. (B) The preferred visual experience for OPA-pLPC depicts solid geometric surfaces of buildings and streets together with constant speed, indicating that the OPA-pLPC is most active while the participant moves at steady speed through a geometric man-made environment. The anti-preferred visual experience depicts stationary periods dominated by trees and foliage, whose irregular texture contrasts with the geometric surfaces of the buildings. This pattern suggests that the OPA-pLPC is engaged in representing the geometry of the scene and recruited for navigational actions. (C) The preferred visual experience for PPA-ColS also depicts buildings and constant speed, though the buildings appear more blurred than in OPA-pLPC, indicating that the PPA-ColS is most active while the participant moves past buildings whose appearance changes across frames. The anti-preferred visual experience depicts stationary periods in areas more open than those for OPA-pLPC. This pattern suggests that the PPA- ColS is more sensitive to changing scene appearance than to stable geometric surfaces.

The preferred visual experience for the occipital place area-posterior lateral parietal cortex region (OPA-pLPC) depicts the solid geometric surfaces of buildings and streets together with constant speed (Fig. 3B). The anti-preferred visual experience depicts stationary periods dominated by trees and foliage, whose irregular texture contrasts with the geometric building surfaces seen in the preferred visual experience. OPA-pLPC is therefore most active while the participant moves at steady speed through environments with strong geometric structure such as the city, and least active during stationary periods and in geometrically irregular environments such as rural regions. This pattern agrees with our encoding models, which suggest that OPA-pLPC represents information for visually-guided movements (Persichetti & Dilks, 2018). The preferred visual experience indicates that OPA-pLPC represents the geometric structure of the scene and that it is relatively invariant to texture, a useful property for action planning. The preference for dynamic over static scenes further indicates that the activity of OPA-pLPC is tied to concrete navigational actions.

The preferred visual experience for the parahippocampal place area-collateral sulcus region (PPA-ColS) also depicts buildings and constant speed, though the buildings appear more blurred than the preferred visual experience for OPA-pLPC (Fig. 3C). The anti-preferred visual experience depicts stationary periods in areas that are more open than the anti-preferred visual experience for OPA-pLPC. PPA-ColS is therefore most active while the participant moves at steady speed past different structures, and least active during stationary periods in open areas that offer little change in scene content. This pattern agrees with our encoding models, which suggest that PPA-ColS represents semantic and identity information. The preferred visual experience indicates that PPA-ColS is more sensitive to the changing appearance of a scene than it is to stable geometric surfaces, and that its representation may be more semantic than the geometric representation in OPA-pLPC.

### Preferred visual experiences for parietal navigation regions

We next examined three regions in the parietal cortex, the anterior lateral parietal cortex region (aLPC), the dorsal precuneus region (dPrCu), and the anterior precuneus region (aPrCu). The preferred visual experience for the anterior lateral parietal cortex region (aLPC) includes a weak “go to” prompt, buildings, and acceleration across the frames (Fig. 4A). The anti-preferred visual experience depicts a stationary period with foliage and another vehicle ahead of the participant. Thus, aLPC is most active as the participant begins a trial and moves through a geometric man-made environment, and least active when the participant is stopped behind another vehicle. This pattern builds upon our encoding models, which suggest that aLPC supports the production of visually-guided navigational actions and the integration of concrete actions with abstract plans. The weak presence of the “go to” prompt suggests that aLPC, like RSC-pPrCu, may be engaged by route planning at the beginning of trials.

**Figure 4.**
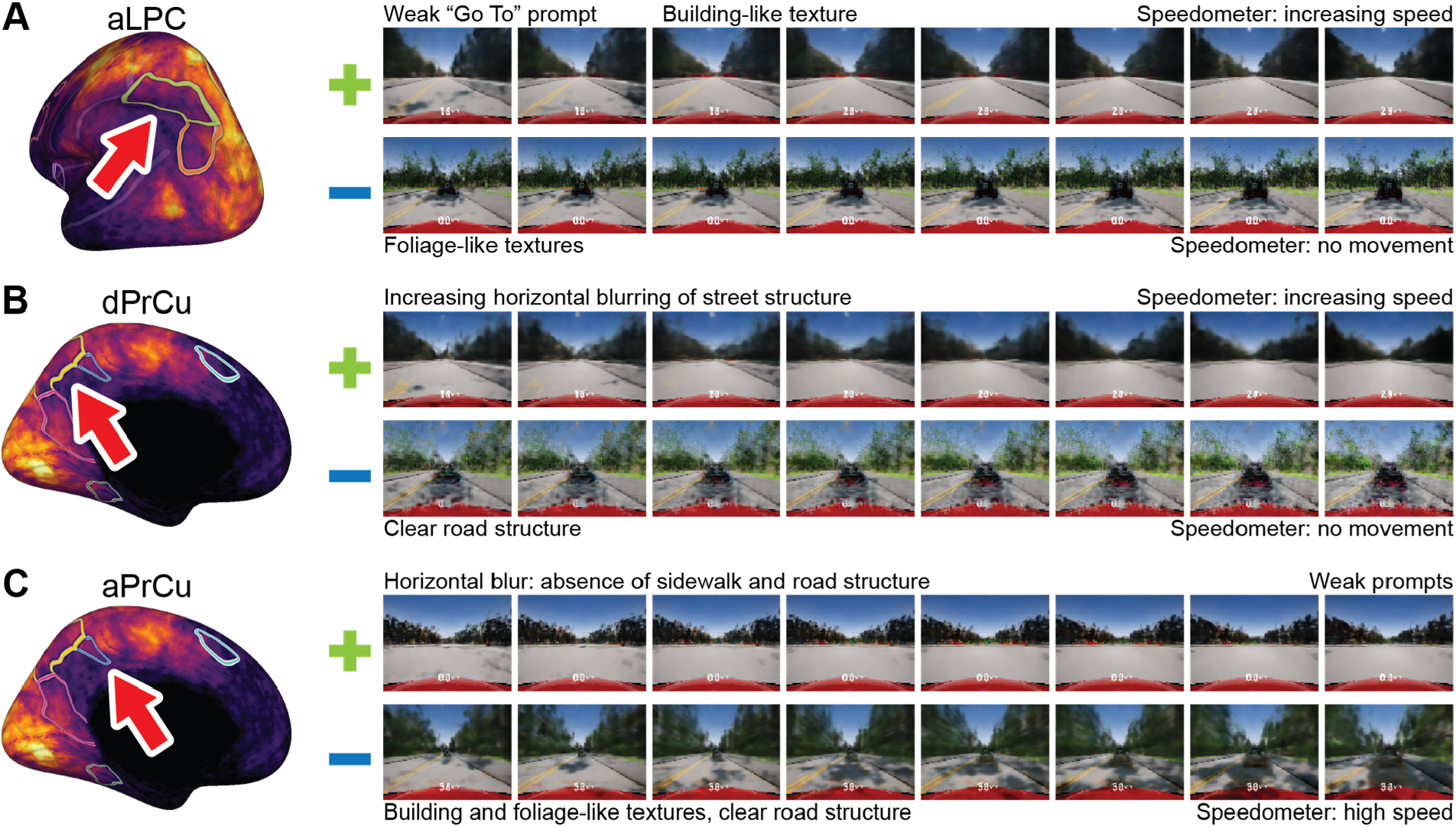
Preferred and anti-preferred visual experiences for parietal navigation regions. Our prior study of the navigation network revealed three regions in the parietal cortex, and suggested that these regions perform sensorimotor transformations to produce concrete navigation actions. (A) The preferred visual experience for aLPC includes a weak “go to” prompt, buildings, and acceleration over the frames (visible in the reconstructed speedometer values at the bottom of each frame), indicating that the aLPC is most active as the participant begins a trial and when moving through a geometric man-made environment. The anti-preferred visual experience depicts a stationary period with foliage and another vehicle in front of the participant. This pattern suggests that the aLPC may be engaged by route planning at the beginning of trials. (B) The preferred visual experience for dPrCu depicts acceleration over the frames and a blurring of the sidewalk and road-line structure suggestive of turns, indicating that the dPrCu is most active when the participant accelerates and steers through turns. The anti-preferred visual experience depicts a stationary period. This pattern suggests that the dPrCu is active when participants execute concrete navigational actions. (C) The preferred visual experience for aPrCu shows an absence of road lines and sidewalks, suggestive of intersections and open navigation affordances, and weak presence of prompts, indicating that the aPrCu is most active when multiple routes are available, and at the start and end of trials. The anti-preferred visual experience shows strong road structure at constant speed. This pattern suggests that the aPrCu is involved in representing not only navigational affordances, but also goals.

The preferred visual experience for the dorsal precuneus region (dPrCu) depicts acceleration across the frames, and a blurring of the sidewalk and road line structure suggestive of turns (Fig. 4B). The anti-preferred visual experience depicts a stationary period behind another vehicle, and also foliage-like textures. Thus, the dPrCu is most active when the participant accelerates and steers through turns, and least active when the participant is stopped. This pattern agrees with our encoding models, which suggest that dPrCu represents information for planning movements through the visual scene. The preferred visual experience indicates that dPrCu is active when participants execute concrete navigation-related actions, such as turns.

The preferred visual experience for the anterior precuneus region (aPrCu) shows an absence of road lines and sidewalks consistent with the visual structure of intersections, low speed, and weak evidence for the “go to” and “arrived” prompts (Fig. 4C). The anti-preferred visual experience shows strong road structure at constant speed. Thus, the aPrCu is most active as the participant slows at intersections, where multiple routes are available, and at the start and end of trials, and least active while cruising along a well-defined road. This pattern expands upon our encoding models, which suggest that aPrCu represents information for planning concrete navigational actions. The preferred visual experience indicates that aPrCu may be involved both in representing the navigational affordances that are critical to action planning and in representing navigational goals.

### Preferred visual experiences for prefrontal navigation regions

Finally, we examined five prefrontal regions that support navigation, namely the posterior medial prefrontal cortex region (pmPFC), the posterior superior frontal sulcus region (pSFS), the inferior frontal sulcus region (IFS), the frontal pole region (FP), and the anterior insula region (aIns). The preferred visual experience for pmPFC shows buildings and deceleration toward a stop (Fig. 5A). The anti-preferred visual experience shows acceleration, foliage-like patterns, and the appearance of road lines in later frames. Thus, the pmPFC is most active as the participant slows, possibly as they approach a destination, and least active when exiting a turn.

**Figure 5.**
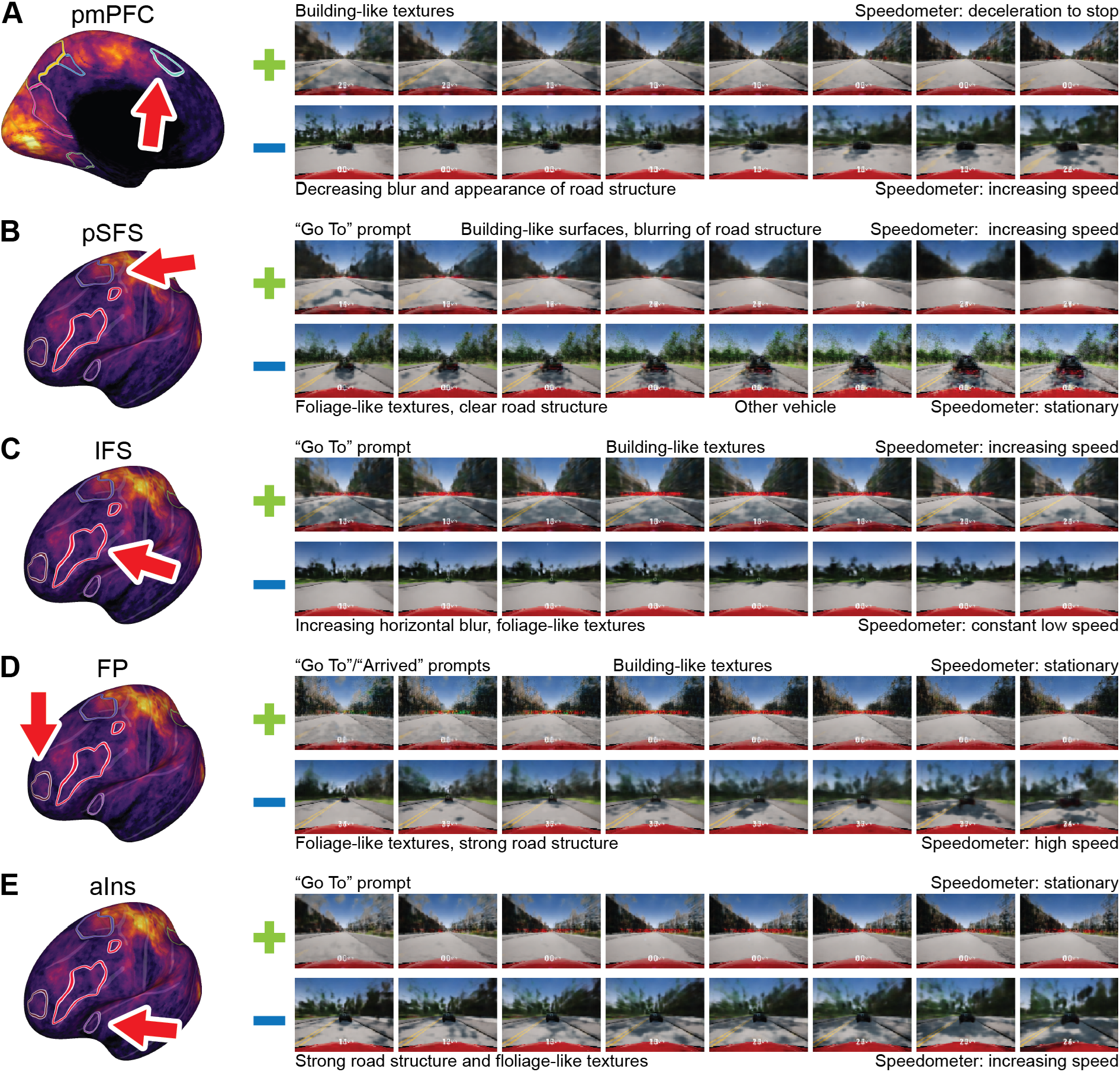
Preferred and anti-preferred visual experiences for prefrontal navigation regions. Our prior study of the navigation network revealed five regions in the prefrontal cortex, and suggested that these regions represented the most abstract aspects of spatial navigation. (A) The preferred visual experience for pmPFC shows buildings and deceleration toward a stop, indicating that the pmPFC is most active as the participant slows, likely as they approach a destination. The anti-preferred visual experience shows acceleration, foliage- like patterns, and road lines suggestive of exiting a turn. This pattern suggests that the pmPFC may be involved in identifying the destination and anticipating arrival. (B) The preferred visual experience for the pSFS shows the “go to” prompt, acceleration, building-like structures, and a disappearance of road lines toward the end of the frames suggestive of turns, indicating that the pSFS is most active at the beginning of trials, and during increasingly rapid movement through the environment. The anti-preferred visual experience depicts a stationary period behind another vehicle amid foliage-like textures. This pattern suggests that the pSFS is an integration region for many different navigation-related representations. (C) The preferred visual experience for the IFS shows a strong “go to” prompt, acceleration over the frames, and building-like textures, indicating that the IFS is most active at the beginning of trials, and when moving through a dense urban environment. The anti- preferred visual experience depicts low-speed turns with blurred road structures on a foliage background. This pattern suggests that the IFS may be involved in route planning at the beginning of trials and in responding to changing scene identities. (D) The preferred visual experience for the FP shows both the “go to” and “arrived” prompts and a noisy building-like texture, indicating that the FP is most active at both the start and end of trials. The anti-preferred visual experience depicts strong road structure and cruising at constant speed. This pattern suggests that the FP manages abstract navigation plans, but may be less involved in the concrete perception and action aspects of navigation. (E) The preferred visual experience for the anterior insula (aIns) shows the “go to” prompt along with some building-like textures in a stationary period, indicating that the aIns is most active while the participant awaits the beginning of trials and in dense urban environments. The anti-preferred visual experience shows acceleration with strong road structures and foliage-like textures. Unlike the preferred visual experiences for other regions, the “go to” prompt appears only from the second frame onward, suggesting that aIns activity anticipates the prompt. This pattern suggests that the aIns may play a role in anticipating upcoming navigational demands.

This pattern expands upon our encoding models, which suggest that pmPFC uses visual information to update progression along routes. The contrast between decelerating toward buildings and accelerating away from them suggests that pmPFC may be involved in identifying the destination and anticipating arrival.

The preferred visual experience for the posterior superior frontal sulcus region (pSFS) shows the “go to” prompt, acceleration, building-like structures, and a disappearance of road lines toward the end of the frames suggestive of entering turns (Fig. 5B). The anti-preferred visual experience depicts a stationary period behind another vehicle and foliage-like textures. Thus, the pSFS is most active at the beginning of trials when the participants must plan the route, during increasingly rapid movement, and during turns through the environment, and it is least active during stationary periods. This pattern agrees with our encoding models, which suggest that pSFS integrates many aspects of dynamic spatial navigation. The range of features in its preferred visual experience, spanning the “go to” prompt, motion, building structures, and turns, suggests that pSFS acts as an integration center that represents many types of information as participants actively navigate.

The preferred visual experience for the inferior frontal sulcus region (IFS) shows a strong “go to” prompt, acceleration across the frames, and building-like textures (Fig. 5C). The anti-preferred visual experience depicts low-speed turns with blurred road structure on a foliage background. The IFS is therefore most active at the beginning of trials when participants are most likely engaged in route planning, and when moving rapidly through a dense urban environment, and it is least active during slow turns and also in more rural environments. This pattern expands upon our encoding model results, which suggest that IFS supports attentional control for navigation. The preferred visual experience suggests that IFS may be engaged in route planning at the beginning of trials, and may also respond to changing scene identity.

The preferred visual experience for the frontal pole region (FP) shows both the “go to” and “arrived” prompts and a noisy texture that resembles buildings (Fig. 5D). The anti-preferred visual experience depicts strong road structure and cruising at constant speed. FP is therefore most active at both the start and end of trials, when navigational goals are set and met, and least active during the steady driving that occurs between them. This pattern builds upon our encoding models, which indicate that FP is a convergence region for many types of abstract navigation-related representations. The contrast between the preferred and anti-preferred visual experiences suggests that FP manages abstract navigation plans but may be less involved in producing concrete action outputs.

Finally, the preferred visual experience for the anterior insula region (aIns) shows the “go to” prompt along with building-like textures during a stationary period (Fig. 5E). The anti-preferred visual experience shows acceleration with strong road structure and foliage-like textures. Thus, the aIns is most active while the participant awaits the beginning of trials and in dense urban environments, and least active during constant movement and in more rural areas. Unlike the preferred visual experiences for the other prefrontal regions, the “go to” prompt appears only from the second frame onward, which suggests that aIns activity may anticipate the prompt. This pattern expands upon our encoding models, which suggest that the aIns aids in tracking progression along a route, and indicates that aIns may actively anticipate upcoming events in navigational sequences.

These preferred visual experiences align well with results from high-dimensional encoding models of these regions. They complement the complex tuning profiles that the encoding models had produced, and show that many of these regions are most engaged during dynamic experiences that combine world events and participant actions. Together, these preferred visual experiences provide a rich and intuitive description for the functional properties of the cortical navigation network.

### Preferred visual experiences can generate novel hypotheses about brain function

The active navigation task elicited broad activity beyond regions for vision, navigation, and action, and the contribution of other regions to navigation is unclear. Because the VAE is trained in a completely data-driven manner, it could reconstruct preferred visual experiences for any region and may offer novel insights into the functional properties of these regions. To test this possibility, we examined the temporoparietal junction (TPJ). The TPJ is not typically associated with navigation, but our prior data suggest that it appears to be engaged by the driving task. The preferred visual experience for the TPJ depicts a vehicle ahead receding into the distance, while the anti-preferred visual experience depicts driving down an empty road (Fig. 6A). The TPJ is therefore most active when the participant is in the presence of another vehicle, and least active in the absence of other vehicles. This pattern contrasts with those of many navigation regions, in which other vehicles instead appear in the anti-preferred visual experience, and suggests that the TPJ may be involved in processing other navigating agents in the world.

**Figure 6.**
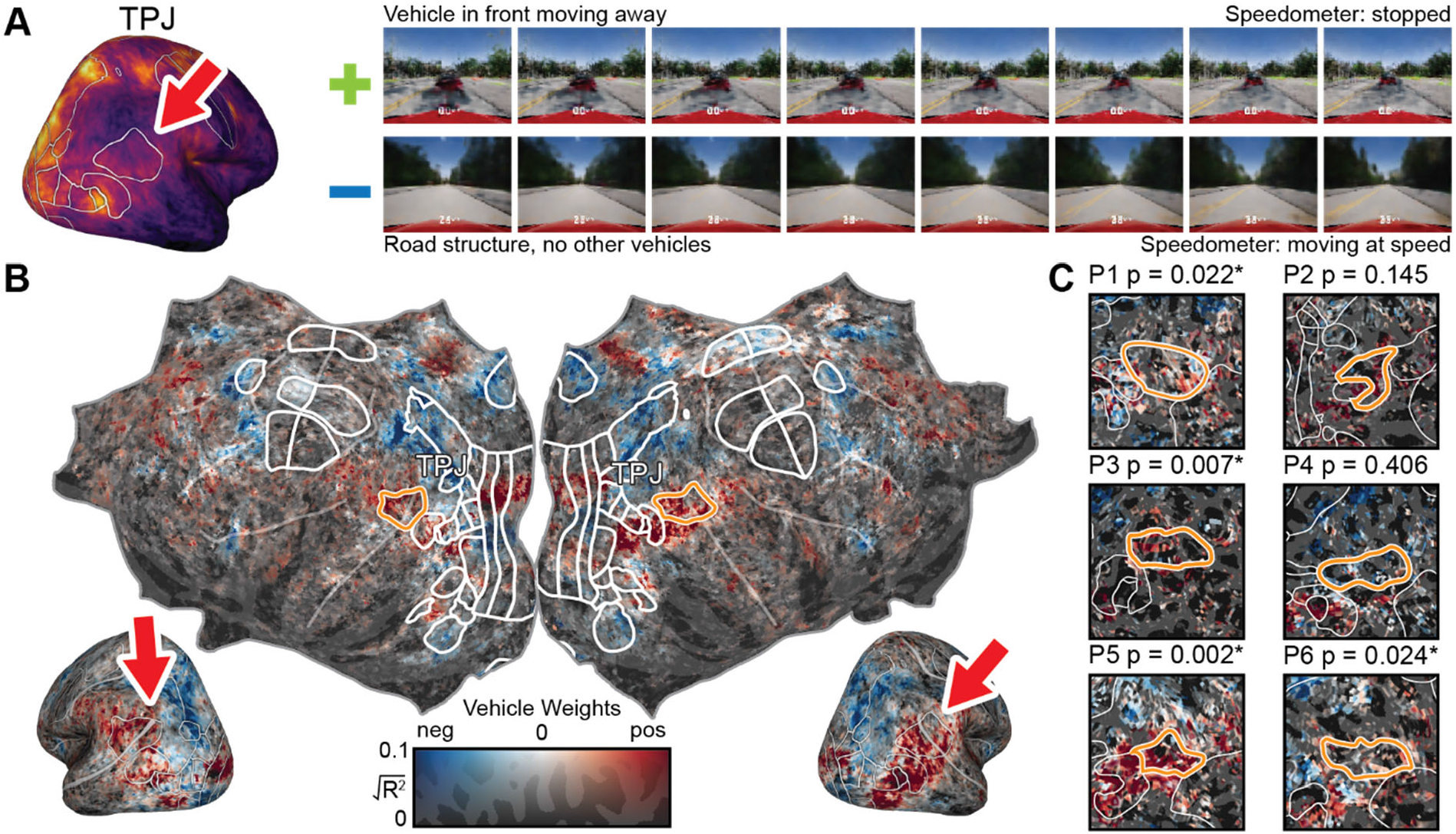
Preferred visual experiences generate novel insights about brain function. Because the VAE latent space is learned from the visual scene alone, it can potentially recover information about any aspect of brain representation that corresponds to structured visual experiences. To explore this possibility, we examined the preferred and anti-preferred visual experiences outside the core navigation network. We focused on the temporoparietal junction (TPJ). Many prior studies have linked the TPJ to theory-of-mind processes, but to our knowledge no studies have examined its role in spatial navigation. (A) The preferred visual experience for TPJ depicts a vehicle ahead receding into the distance, while the anti-preferred visual experience depicts an empty road. The TPJ is therefore most active when the participant is in the presence of another vehicle, and least active in the absence of other vehicles, suggesting that the TPJ may be involved in processing other navigational agents in the world. (B) To corroborate this observation, we examined weights for “vehicles” from an explicit encoding model for visual semantics. Average vehicle semantic weights across participants are shown on flattened and inflated cortical surfaces. Red indicates positive weights and blue indicates negative weights. The TPJ is outlined on the flattened surface and indicated by arrows on the inflated surfaces. The TPJ shows significant weights for vehicles (p = 0.005, permutation test). (C) Closeups of the flattened surface of the right TPJ are shown for six individual participants, using the same color scheme as for panel B (For full maps, see Supplementary Fig. 2). Four of six participants show significant weights for vehicles in the TPJ (p < 0.05, permutation test). Thus, these maps consistently show strong positive weights for vehicles in the TPJ, in agreement with its preferred visual experience. These results suggest that the TPJ may support social cognition related to other vehicles during navigation. This unexpected result demonstrates that VAE-derived preferred visual experiences may provide novel insights about brain function.

To corroborate this observation, we examined the weights for “vehicles” from an explicit encoding model for visual semantics (Zhang & Gallant, 2026). These weights show that the TPJ is significantly selective for vehicles at the group level (Fig. 6B, p = 0.005, permutation test), and this selectivity is found in four of six individual participants (Fig. 6C, p < 0.05, permutation test, and Supplementary Fig. 2). These weights are consistent with the presence of vehicles in the preferred visual experience for the TPJ, suggesting that it may have a role in social cognition for spatial navigation. Previous studies have linked the TPJ to theory-of-mind processes (Saxe & Kanwisher, 2003), but not in the context of spatial navigation. Thus, this finding suggests that VAE-derived preferred visual experiences may provide novel hypotheses about brain function.

## DISCUSSION

In this report, we used the embedding space of a variational autoencoder (Kingma & Welling, 2019) to generate preferred visual experiences for 11 different regions comprising the cortical navigation network (Zhang et al., 2025). In this previous study, high-dimensional encoding models based on hand-crafted feature spaces revealed that functional representations are distributed across these regions, and that each region represents a complex combination of features. Although these regions are functionally distinct, the exact functions are difficult to understand based on the high-dimensional encoding models alone. The VAE-derived preferred visual experiences presented here complement the earlier encoding model analyses of these regions. By mapping the complex tuning properties of each region into an intuitively understandable format in image space, the preferred visual experiences help reveal the functional properties of these regions in more detail. These preferred visual experiences reveal that the regions in the cortical navigation network are most engaged during dynamic combinations of world events and participant actions. Because the VAE is data driven, it can recover the preferred visual experience for any brain region, potentially generating new insights into brain function.

The dynamic nature of these preferred visual experiences is a marked advancement over previous reconstruction efforts that have largely focused on static images (Horikawa & Kamitani, 2017; Lin et al., 2022; Ozcelik & VanRullen, 2023; Ren et al., 2021; Shen et al., 2019). Because the VAE used in this study was trained on 2- second video clips, it learned the short-timescale visual dynamics of navigation, such as optic flow and changes in vehicle speed. The preferred visual experiences consistently contain a legible speedometer that changes smoothly, indicating that the VAE decodes coherent dynamics rather than arbitrary sequences of visual scenes. Notably, the VAE does not always decode sequences that depict motion, as evidenced by many of the anti-preferred visual experiences that depict stationary periods. Thus, egomotion in the preferred visual experiences likely reflects the tuning of these regions rather than a bias in our VAE towards dynamic output. Our results suggest that many regions in the navigation network prefer dynamic scenes, and thus dynamic stimuli and interactive tasks (Cisek & Kalaska, 2010; Huk et al., 2018) are necessary for fully characterizing the navigation network in the human cerebral cortex.

Because the VAE generates a preferred visual experience for every brain region, it can be used to gain new insights about heretofore unknown functions of well-studied brain areas. For example, prior studies that used narrative-based false belief (Saxe & Kanwisher, 2003) or perspective-taking (Saxe & Wexler, 2005) tasks suggest that the TPJ plays an important role in social cognition (Saxe & Kanwisher, 2003). Here, we find that the TPJ also appears to represent aspects of social cognition that are important for navigation in multi-agent environments. This result suggests that new experiments that specifically target social interactions during spatial navigation (Danjo et al., 2018; Omer et al., 2018) may reveal novel functional properties of the brain.

The preferred visual experiences developed here only recover the visual portion of the perception-cognition-action loop that underpins active navigation (Cisek & Kalaska, 2010; Land & Lee, 1994). The participants’ plans and actions that shape the scene are not directly reconstructed; rather, the preferred visual experiences contain their visual correlates, such as blurred street structures and appearance of prompts. Our interpretations about planning or action therefore rest on inference from these correlates, rather than direct readouts.

Furthermore, although complex tasks such as driving broadly engage many functional networks in the brain, the VAE cannot extrapolate beyond the domain of the data on which it was trained (Shirakawa et al., 2025; Udandarao et al., 2024). The preferred visual experiences cannot reflect the full functional properties of each region, but only those associated with the specific task and stimuli on which the VAE was trained. Therefore, the preferred visual experiences reconstructed here are highly biased toward the driving task. It is likely that the functional representations examined here also underpin other behavior in other navigation contexts, such as walking through the environment or navigating indoors.

The preferred visual experiences examined here complement the high-dimensional encoding models fit in our previous study on spatial navigation (Zhang et al., 2025). This interpretive method is general and can be applied to any experiment by training a VAE on its stimulus space, and then fitting an encoding model with the embedding space of that VAE. However, for the preferred visual experiences to be informative, this approach is most well-suited for naturalistic experiments with rich stimuli (Leopold & Park, 2020; Matusz et al., 2019; Nastase et al., 2020; Wu et al., 2006). Rich, complex stimuli that broadly engage the brain are also crucial for data-driven discoveries outside the target regions, such as the example in the TPJ. This combination of high-dimensional encoding models and intuitive preferred visual experiences will become increasingly useful as cognitive neuroscience adopts dynamic tasks to understand complex brain functions under naturalistic conditions.

## METHODS

### Task

The navigation data were collected in (Zhang et al., 2025). Briefly, six participants (3 male, 3 female, ages 24- 34) performed a taxi-driver task in a virtual city while brain activity was recorded with fMRI. A large dynamic virtual city was constructed with Unreal Engine 4 and CARLA (Dosovitskiy et al., 2017), and was populated with both vehicular and pedestrian traffic. Participants used a custom MR-compatible steering wheel and pedal set to drive a car through the virtual world from a first-person perspective. Prior to scanning, participants learned the layout of the world. In the scanner, the taxi-driver task was divided into trials. On each trial, a participant was prompted to navigate to a particular location in the world by instructions displayed on the screen; this prompt is the “go to” prompt. Participants were instructed to use their knowledge of the world to drive to the prompted location via the quickest path, while respecting all traffic rules. After the participant arrived at the destination, another prompt was displayed to acknowledge that they had arrived; this prompt is the “arrived prompt.” A new trial began after a jittered intertrial interval. Participants were allowed to freely view the screen while driving, and eyetracking data were collected at 60 Hz. Trials were recorded with Unreal Engine’s built-in demo recording system. Data were collected in 11-minute runs. 120 minutes of data were collected from participant 1 in two sessions, and 180 minutes of data each were collected from participants 2-6 in three sessions.

The experimental procedures were approved by the Institutional Review Board at the University of California, Berkeley, and written informed consent was obtained from all participants.

### fMRI data collection and processing

MRI data were acquired on a 3T Siemens Trio with a 32-channel head coil, located at the University of California, Berkeley. BOLD data were acquired using a T2*-weighted gradient-echo EPI sequence customized with a water-excitation radiofrequency pulse to prevent contamination from fat signal (TR = 2.0045 s, echo time = 34 ms, flip angle = 74°, voxel size = 2.24 × 2.24 × 3.5 mm^3^, field of view = 224 × 224 mm^2^, matrix size = 100 × 100, and 30 axial slices to cover the entire cortex). Custom personalized headcases were used to stabilize the head and to reduce motion artifacts (Power et al., 2019). Before each functional run, a GRE fieldmap was collected for distortion correction. To facilitate cortical surface reconstruction, anatomical data were also collected (three-dimensional T1-weighted MP-RAGE sequence, 1 × 1 × 1 mm^3^ voxel size and 256 × 212 × 256 mm^3^ field of view).

Respiration and heart rate were recorded using a BIOPAC MP150 system (BIOPAC Systems, Inc.). Eyetracking data were collected using an Avotec dark-pupil IR eyetracker at 60 Hz. The video data were processed with custom software to extract gaze locations (https://github.com/gallantlab/Eyetracking). To ensure accurate eyetracking calibration, at the beginning of every 11-minute functional run, 35 calibration points were presented for 2 seconds each (accuracy: 1.2 ± 0.3 degrees of visual angle, mean ± std. across participants). The taxi-driver task began immediately after the eye calibration sequence was completed.

### fMRI data preprocessing

Each functional run was first motion-corrected using the FMRIB Linear Image Registration Tool (FLIRT) from FSL 5.0 (Jenkinson et al., 2002; Jenkinson & Smith, 2001). Next, functional images were unwarped by applying FUGUE from FSL to fieldmaps collected between functional runs. All volumes in the run were then averaged across time to obtain a high-quality template volume. To align data collected across multiple sessions and runs, the template volume from the first session was selected as a target, and the template volume from all other runs across all sessions were aligned to this target. Pycortex (Gao et al., 2015) was then used to align the functional runs to the anatomical surface. Alignment was checked manually and adjusted as necessary to improve accuracy. Low-frequency voxel response drift was identified using COMPCOR (Behzadi et al., 2007) and removed from the signal. Physiological signals from respiration and heartbeats were regressed out with RETROICOR (Glover et al., 2000). Voxel activity in each 11-minute run was z-scored separately; that is, within each run, the mean response for each voxel was subtracted and the remaining response was scaled to have unit variance. To remove confounds from the eyetracking calibration sequence and detrending artifacts, the first 35 and last 5 TRs were then discarded from each run.

### Cortical surface reconstruction

Freesurfer (Dale et al., 1999) was used to reconstruct cortical surface meshes from the T1-weighted anatomical volumes. The freesurfer anatomical segmentation was checked by hand, and Blender (Blender Foundation) and pycortex were used to manually correct the segmentation where necessary. Standard functional regions were identified with separate localizer experiments (see Supplementary Methods). Blender and pycortex were then used to remove the medial wall, and relaxation cuts were then made into each surface. The cut at the calcarine sulcus was made using retinotopic localizers as a guide to bisect V1 along the horizontal meridian.

### Reconstructing preferred visual experiences

To reconstruct the preferred visual experiences for functional brain regions, we used a variational autoencoder (VAE; Kingma & Welling, 2019) to learn a low-dimensional latent space for the stimulus and task from the screen recordings. The screen recordings were first downsampled from 1024×768 px at 30 fps to 120×90 px at 3.75 fps. In the experiment, all prompts were displayed in white text. Because we wanted to differentiate the prompt types in the pixel values, the stimulus videos were preprocessed to color the “go to” prompts in red and the “arrived” prompts in green.

The VAE used two 3D convolutional neural networks with residual connections as the encoder and decoder, connected by a size-512 bottleneck (see Supplementary Table 1 and Supplementary methods for a detailed description of the architecture). The VAE then was trained on 8-frame windows from the downsampled screen recordings, and optimized with an evidence lower bound objective function. A Gaussian prior with unit diagonal covariance was imposed on the latent space to regularize the features. In particular, the Kullback-Leibler (KL) divergence between the prior and the posterior in the latent space is minimized along with the video reconstruction loss (Kingma & Welling, 2019). All stimulus videos were then mapped to the 512- dimensional latent space of the trained VAE.

Ridge regression was then used to fit a voxelwise encoding model with these latent VAE features. The VAE model weights for each of the six participants were averaged on the fsaverage6 surface. The average VAE model weight was then computed for each of the functional regions in the cortical navigation network (Zhang et al., 2025). Because the ridge parameter affects the scale of the weights, the raw weights were out of the distribution of the embedding vectors that the decoder had been trained on (Supplementary Figs. 3 and 4).

Directly using these raw weights to reconstruct preferred and anti-preferred visual experiences consistently produces scenes near to the average visual experience (Supplementary Figs. 5 and 6). To correct for this scaling issue, the weights were first normalized to unit norm, and then scaled by 10. These rescaled VAE model weights were given to the decoder to reconstruct the preferred visual experience for each region. To produce the anti-preferred visual experience, which corresponds to the lowest activity in each region, the unit norm weights were scaled by -10, and then also given to the decoder. The VAE encoding model was fit separately from the other encoding models so that it had access to all the variance when generating preferred visual experiences.

### Explicit encoding models for visual semantics

Explicit encoding models were built for visual semantics using the game engine’s semantic segmentation (Zhang & Gallant, 2026). To compute visual-semantic features, the game engine was used to semantically segment the participant’s view and determine the category of the object rendered at each pixel. This semantic segmentation covered 16 categories of 3D assets used to create the virtual environment: buildings, pedestrians, vehicles, roads, road lines, sidewalks, traffic signs, poles, walls, fences, fields, ground, foliage, self, miscellaneous objects, and sky. Visual-semantic categories were also included for on-screen display items that were overlaid on the screen rather than rendered in-world: the “go to” and “arrived” cues and the speedometer. The visual-semantic feature space also included 14 additional categories for objects and on-screen display items that appeared during the learning phase, but not during data collection.

Participants are unlikely to allocate equal attention to all areas of the screen, so the visual semantics of the entire screen may not accurately reflect representations of visual semantics under the influence of top-down attention. Therefore, the gaze location was used as a proxy for how participants allocated attention. For each sample in the eyetracking data, a circle of 5° diameter was drawn around the gaze position on the corresponding semantic segmentation frame. The fraction of this circle occupied by each of the 16 categories was then calculated and used as features.

The visual-semantic content of the scene co-varies with many other features, such as the low-level visual structure of the scene, the participants’ motor actions, and other navigation-related parameters. The contributions of these other features to brain activity were captured by 37 additional feature spaces (see Supplementary Table 2 for a list, and (Zhang et al., 2025) for detailed descriptions of each feature space). In total, these 37 feature spaces encompassed 28,101 additional features.

Banded ridge regression (Dupré la Tour et al., 2022; Nunez-Elizalde et al., 2019) was used to fit voxelwise encoding models for the visual semantics features simultaneously with 37 other feature spaces. To capture the hemodynamic response, a finite impulse response filter (FIR) was implemented by including delayed copies of all features at 5 temporal delays. The optimal regularization parameters and model weights were empirically selected by cross-validation on the training set. Model performance was then quantified by the coefficient of determination (R^2^) between the predicted and actual brain activity on the test set. For each voxel, the joint model performance for all models was then partitioned over models for each of the feature spaces. Here we call these values the split R^2^ scores.

To recover the visual semantic tuning weights, the fit visual semantic model weights were averaged across the five FIR filter delays. Because each voxel is regularized independently, the scale of the raw model weights differs across voxels. To account for this scaling difference, these weight vectors were first normalized to have unit norm, and then scaled by the split R^2^ score of the visual semantic model. To create the group-average weights, weights from individual participants were projected to the fsaverage6 surface and then averaged at each vertex.

A permutation test was used to test whether the TPJ was significantly tuned to vehicles. Null weights were obtained by randomly selecting spatially contiguous patches that contained an equal number of vertices to the TPJ. This process was repeated 1000 times in each hemisphere. The average vehicle weight in the left and right TPJ were then compared against their respective nulls. This process was performed at both the group level and within individual participants.

### Code availability

Associated code will be made publicly available on Github upon publication.

## Acknowledgments

We thank all members of the Gallant Lab for discussion and support throughout this work. We also thank Dimitar Filev and Ken Washington at the Ford Motor Company, and Marc Steinberg at the Office of Naval Research, who provided critical funding support during the early stages of this project.

## Funding

National Institutes of Health grant R01-EY031455 (JLG)

Office of Naval Research grant N000142012002 (JLG)

Ford University Research Program (JLG)

Office of Naval Research DURIP awards N00014-22-1-2217 (JLG)

Office of Naval Research Multidisciplinary University Research Initiative grant N000141410671 (JLG)

Keck Foundation grant A25-0064-S001 (JLG)

National Science Foundation Graduate Research Fellowship Program DGE 1106400 & DGE 1752814 (TZ)

## Author contributions

Conceptualization: TZ, CC, JLG

Methodology: CC, TZ, JLG

Investigation: CC

Formal Analysis: CC

Writing – original draft: TZ

Writing – review & editing: TZ, CC, JLG

