## Supplementary Materials for "Preferred visual experiences provide intuitive descriptions of the functional properties of the cortical navigation network"

**The file includes:**

Supplementary figures 1-6

Supplementary tables 1-2

Supplementary methods

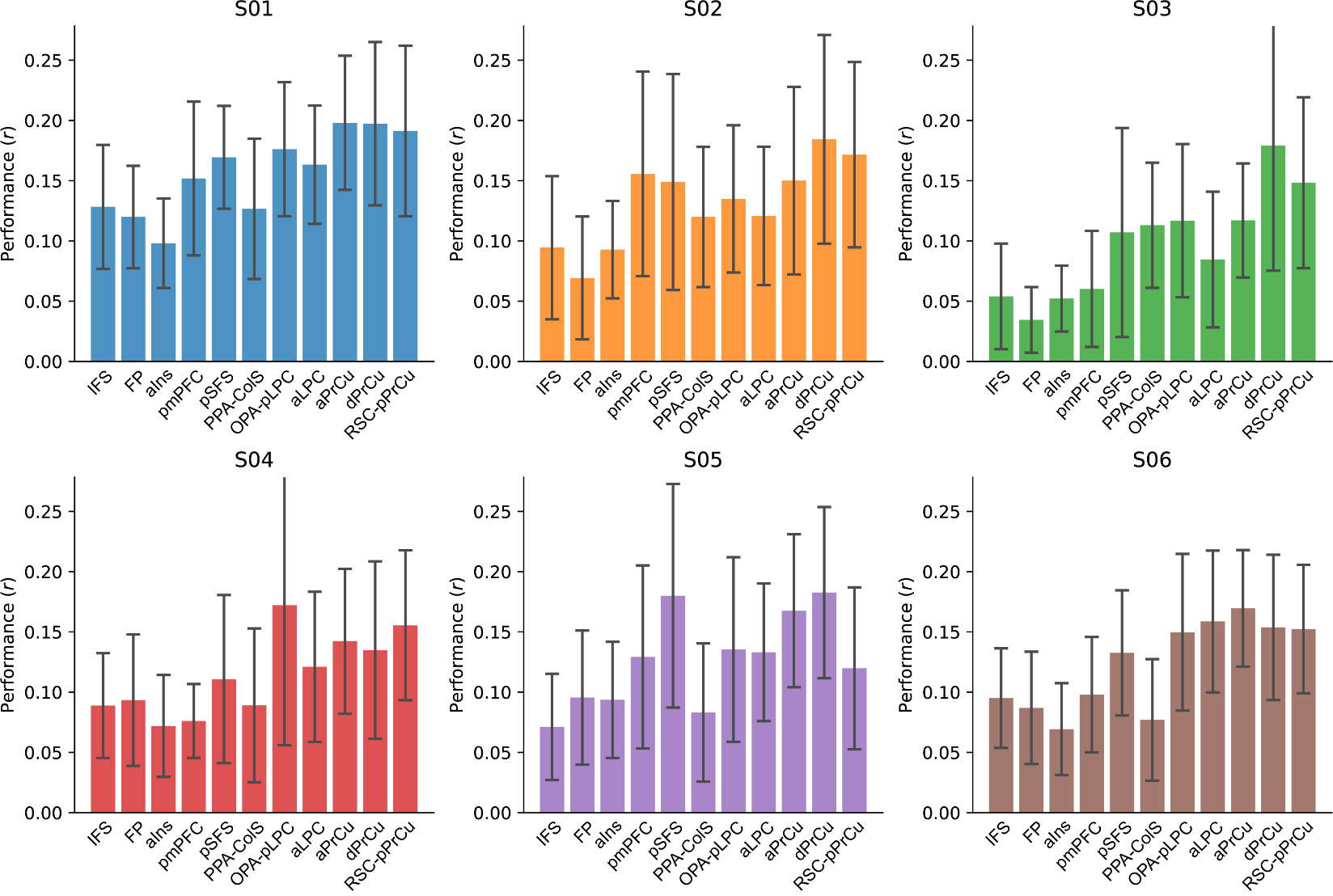

**Supplementary Figure 1. VAE encoding model performance by region in the navigation network.** Here we show voxelwise encoding model performance in each of the 11 regions in the navigation network for each participant. Bars indicate means in each region, and error bars indicate standard deviation.

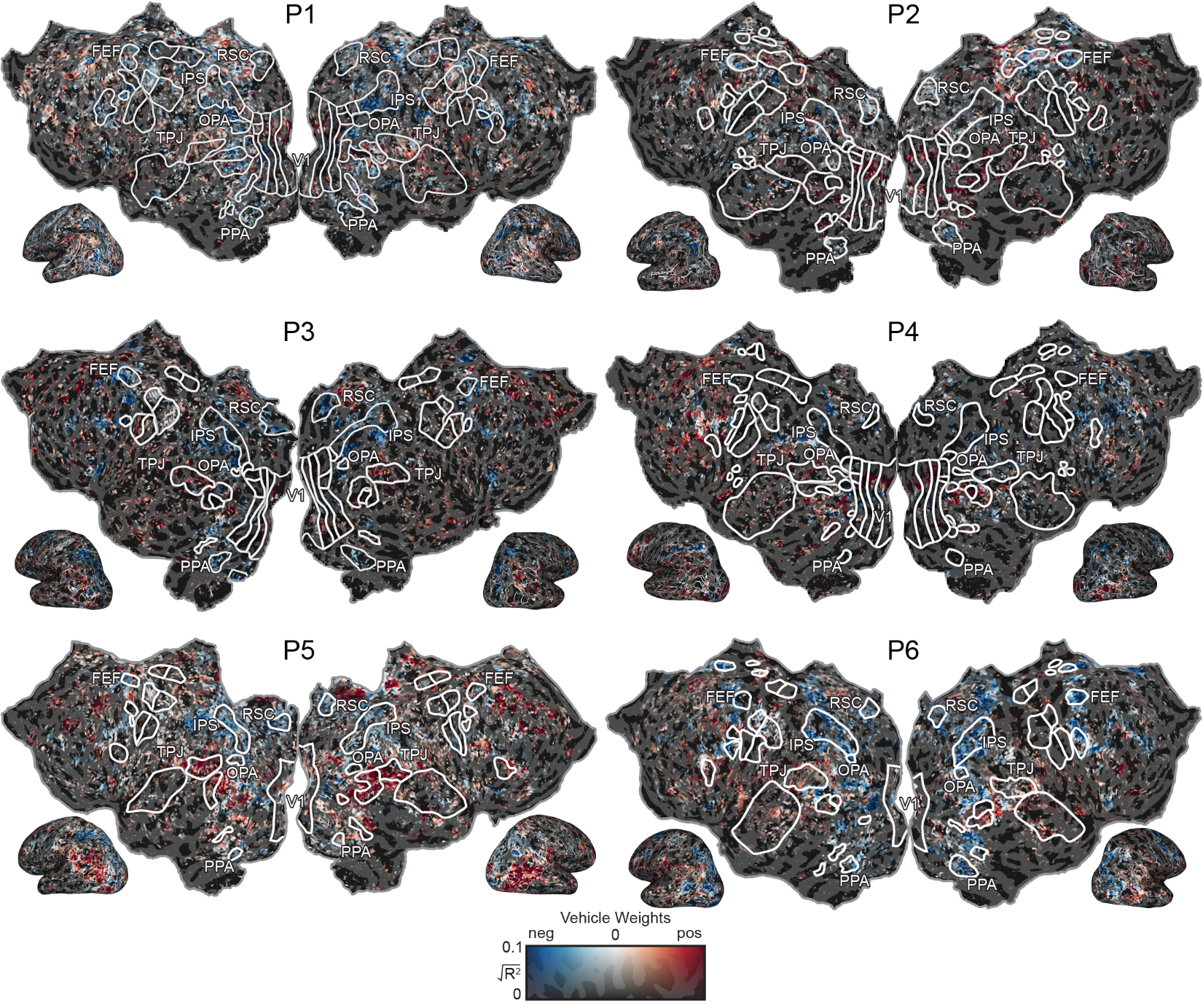

**Supplementary Figure 2. Whole-cortex vehicle weights in individual participants.** Whole-cortex maps of vehicle weights in individual participants. Colors are the same as in Figure 6.

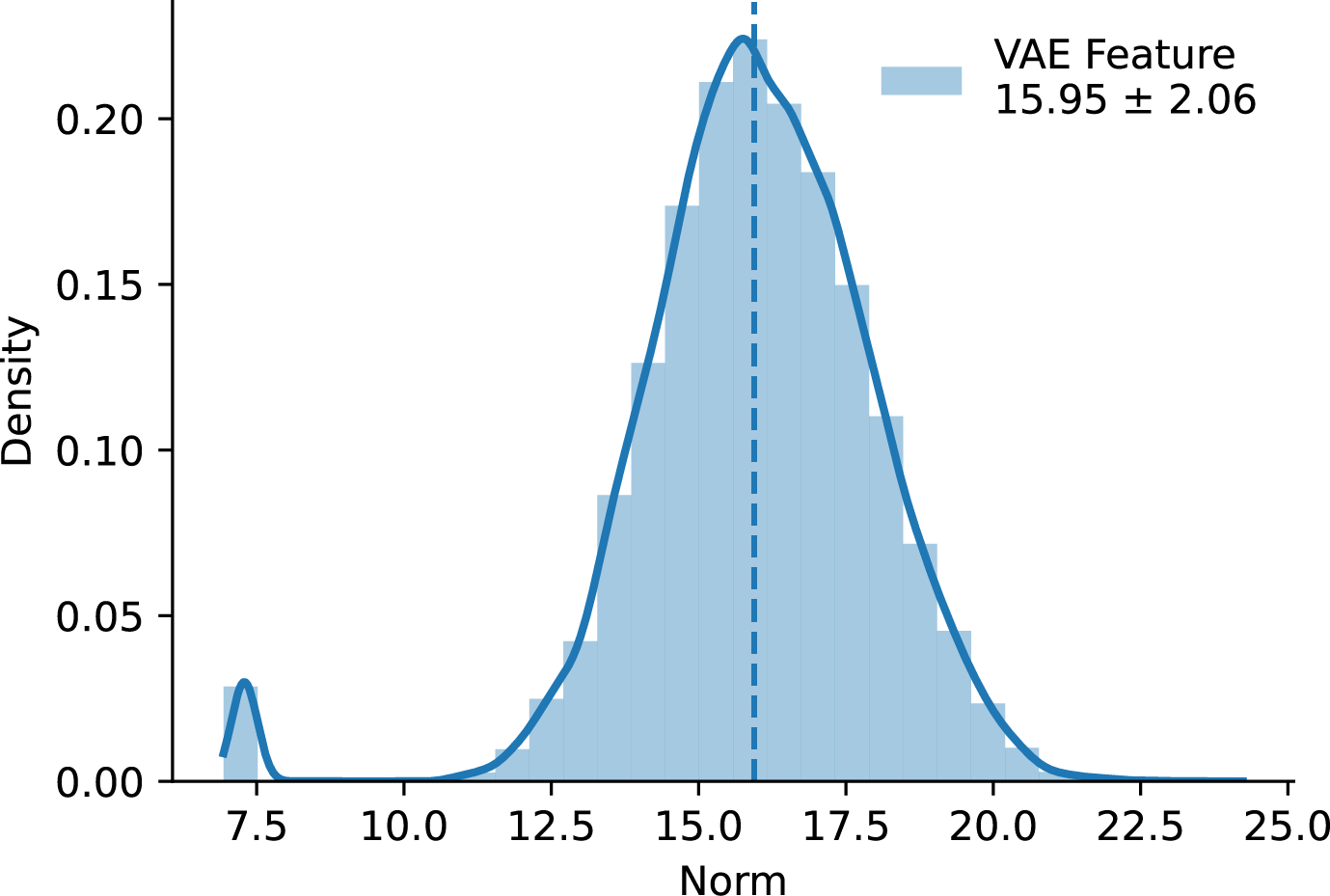

**Supplementary Figure 3. VAE encoding model feature norms.** This figure shows the distribution of norms for the VAE latent embedding features used for fitting voxelwise encoding models. The mean norm of these features is 15.95 and the standard deviation is 2.06.

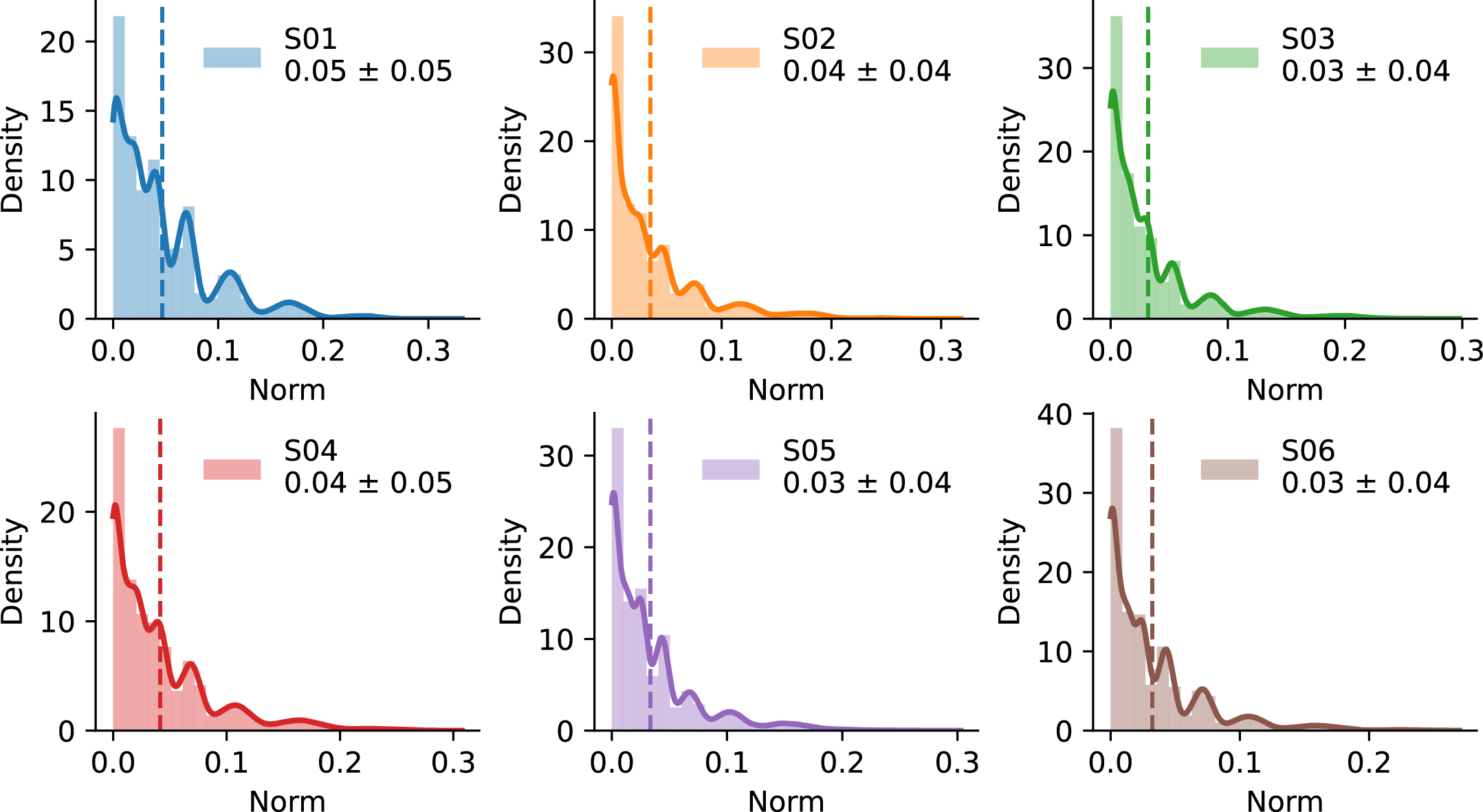

**Supplementary Figure 4. VAE encoding model weight magnitudes differ from the VAE embedding feature magnitudes.** The scale of the voxelwise encoding model weights were largely determined by the regularization parameter. Because each voxel was regularized separately, raw weights across voxels differ in scale and are not directly comparable. To make the weights comparable across voxels, they were rescaled such that their norms corresponded to model prediction performance. Here we show the density histograms of these VAE encoding model weights for each participant. The mean weight norms are on the order of 10^-2^, three orders of magnitude smaller than the VAE feature norms (Supplementary Figure 3). Because the VAE decoder is sensitive to the magnitude of the latent embedding, this discrepancy in magnitudes necessitated rescaling the encoding model weights before they were passed to the decoder.

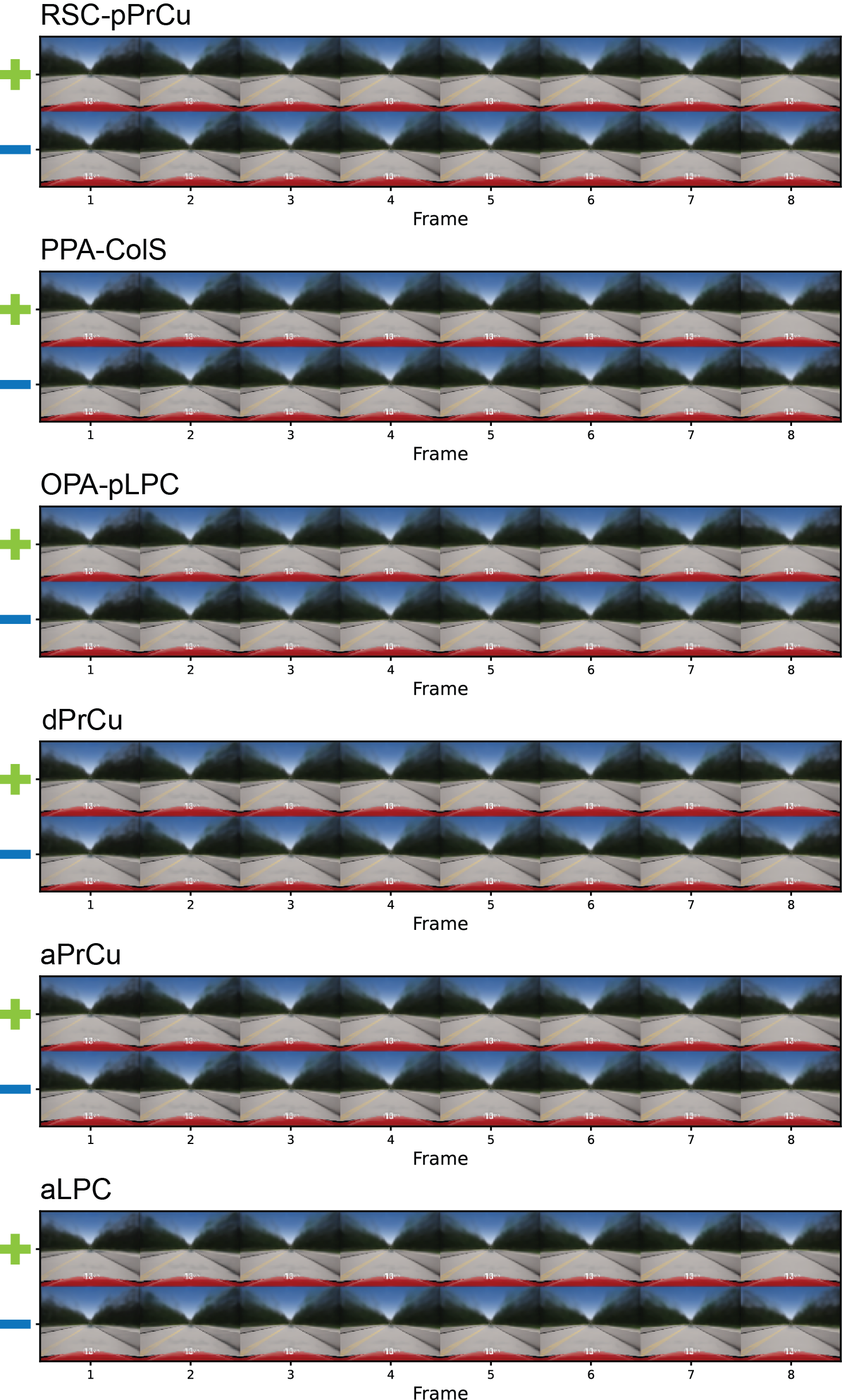

**Supplementary Figure 5. Reconstructed visual experiences without weight rescaling for anterior visual and parietal regions in the navigation network.** Reconstructed visual experiences follow the same format as Figures 2-5. Preferred and anti-preferred visual experiences are reconstructed using the non-rescaled encoding model weights. Because the norm of the encoding model weights were four orders of magnitude smaller than the VAE embedding vectors, directly passing these model weights to the decoder produces visual experiences that are near the average visual experience. For interpretable decoded visual experiences, the weights require rescaling.

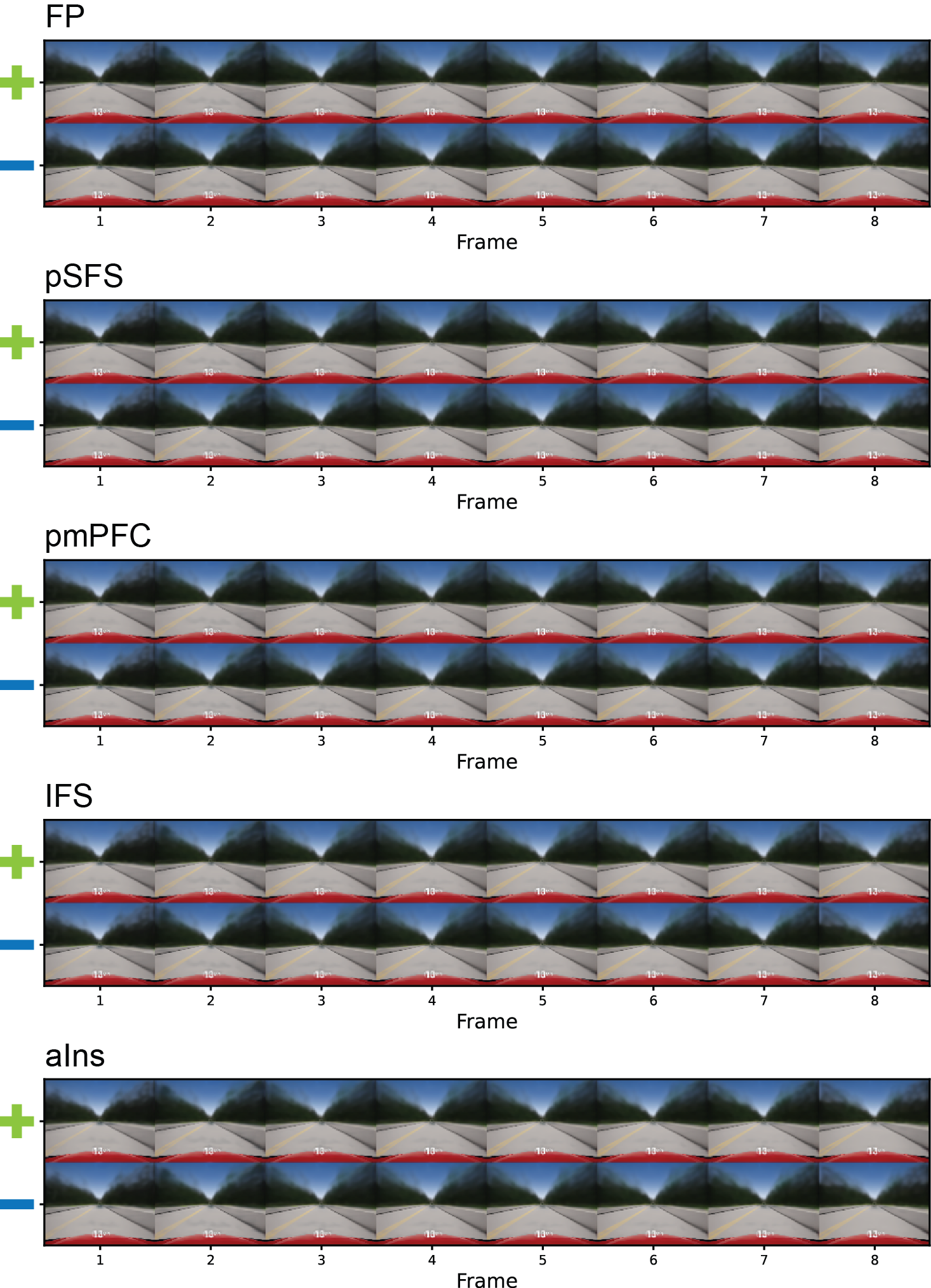

**Supplementary Figure 6. Reconstructed visual experiences without weight rescaling for prefrontal regions in the navigation network.** Reconstructed visual experiences follow the same format as Figures 2-5. Preferred and anti-preferred visual experiences are reconstructed using the non-rescaled encoding model weights.

**Supplementary Table 1. Model configuration of each layer in VAE**

| **Block** | **Kernel (t,h,w)** | **Stride (t,h,w)** | **Normalization** | **Output shape** |
| --- | --- | --- | --- | --- |
| Input | — | — | — | 3 × 8 × 90 × 120 |
| Encoder layer-1 | 1 × 3 × 3 | 1, 1, 1 | batch norm | 64 × 8 × 90 × 120 |
| Encoder layer-2 | 3 × 3 × 3 | 2, 2, 2 | batch norm | 128 × 4 × 45 × 60 |
| Encoder layer-3 | 3 × 3 × 3 | 2, 2, 2 | batch norm | 256 × 2 × 23 × 30 |
| Encoder layer-4 | 3 × 3 × 3 | 2, 2, 2 | none | 512 × 1 × 12 × 15 |
| Global average pool | — | — | — | 512 |
| Decoder  fully connected, reshaped | — | — | none | 256 × 2 × 23 × 30 |
| Decoder layer-1 | 3 × 3 × 3 | 1, 1, 1 | batch norm | 128 × 4 × 45 × 60 |
| Decoder layer-2 | 3 × 3 × 3 | 1, 1, 1 | batch norm | 64 × 8 × 90 × 120 |
| Output head | 3 × 3 × 3 | 1, 1, 1 | — | 3 × 8 × 90 × 120 |

**Supplementary Table 2**. A brief description of the other 37 feature spaces used in the voxelwise encoding models alongside the visual semantics model for the navigation experiment.

| Feature Space | Short Description |
| --- | --- |
| Eyetracking power | The aggregate power across channels encoding gaze location and its derivatives in screen space encoded in cartesian and polar coordinates |
| Gaze grid | Gaze location in screen space encoded with a hexagonal grid basis (Nau et al., 2018) |
| Framewise motion-energy | Spatiotemporal Gabor energy in screen coordinates (Nishimoto & Gallant, 2011) |
| Retinotopic motion-energy | Spatiotemporal Gabor energy in retinotopic coordinates |
| Frame semantics | Semantic content of the screen |
| Depth | Distance to objects in the virtual world |
| Scene structure | The distribution of visible surfaces and their orientations relative to the participant’s viewpoint (Lescroart & Gallant, 2019) |
| Spatial semantics | Visible semantic content of the virtual world around the participant in egocentric coordinates |
| Navigational affordance | Where in the scene that the participant can move through (Bonner and Epstein, 2017) |
| Controls | Steering, gas, brake, and gear selections from the subject |
| Future path | Future path that the participant will take in egocentric coordinates |
| Distance to next turn | Fraction of total distance travelled until the next turn encoded with a Fourier basis (Nitz, 2006) |
| Time to next turn | Fraction of total time travelled until the next turn encoded with a Fourier basis |
| Beeline distance remaining | Beeline distance remaining to the destination (Howard et al., 2014) |
| Path distance remaining | Distance remaining along the actual planned path to the destination (Howard et al., 2014) |
| Spatial route progression (continuous) | Fraction of total distance travelled to the destination encoded by a Fourier basis (Alexander and Nitz, 2017) |
| Temporal route progression (continuous) | Fraction of total time travelled to the destination encoded by a Fourier basis |
| Spatial route progression (binned) | Fraction of total distance travelled to the destination encoded by a set of indicator features |
| Temporal route progression (binned) | Fraction of total time to the destination encoded by a set of indicator features |
| Destination vector | An egocentric vector to the destination (Howard et al., 2014; Brown et al., 2016; Ormond and O’Keefe, 2022) |
| Destination-anchored vector | An allocentric vector from the destination to the current location of the participant |
| Destination grid representation | Position of the destination in the world encoded with a hexagonal grid basis |
| Allocentric path integration | Allocentric vector to the start location of the current trial (Sherrill et al., 2013; Chrastil et al., 2015) |
| Egocentric path integration | Egocentric vector to the start location of the current trial |
| Beeline distance elapsed | Beeline distance from the start of the current trial (Sherrill et al., 2013; Chrastil et al., 2015) |
| Path distance elapsed | Actual path distance travelled from the start of the current trial |
| Time elapsed | Time elapsed since the start of the current trial encoded by a set of indicator features |
| Time elapsed phase | Time elapsed since the start of the current trial encoded by a Fourier basis |
| Place fields | Place field-like representations of the participant’s 2D location in the virtual world(O’Keefe and Dostrovsky, 1971; Ekstrom et al., 2003) |
| Grid cells | Features with grid cell-like periodic spatial selectivity on a hexagonal grid in the virtual world (Fyhn et al., 2004; Sargolini et al., 2006) |
| Head direction | The head direction of the participant in the world encoded with a set of indicator bins (Taube et al., 1990; Vass and Epstein, 2013) |
| 6-fold Head direction | The head direction of the participant in the world, with 6-fold symmetry, encoded with a quadrature filter (Doeller et al., 2010) |
| Gaze direction | The gaze direction of the participant in the world relative to the current heading direction encoded with a set of indicator bins |
| 6-fold gaze direction | The gaze direction of the participant in the world relative to the current heading direction, with 6-fold symmetry, encoded with a quadrature filter |
| Road graph | The planned path of the subject expressed as a set of edges on a graphical representation of the road network (Chrastil and Warren, 2014; Javadi et al., 2017; Warren et al., 2017) |
| Pedestrians | Positions of pedestrians in egocentric coordinates (Danjo et al., 2018; Omer et al., 2018) |
| Vehicles | Positions of other cars in egocentric coordinates |

**Supplementary Methods**

Variational Autoencoder fitting

*Stimuli preprocessing*

In the experiment, both the “go to <destination>” and “arrived at <destination>” prompts were displayed in white text to the participants. Because the VAE operates in pixel space, it does not inherently differentiate between the two types of prompts. To preserve the identity of the prompts in the VAE embedding, the screen recordings were preprocessed to color the “go to <destination>” prompts red and the “arrived at <destination>” prompts green.

*Stimuli*

The stimulus corpus consisted of first-person driving videos recorded during scanning: 112 driving runs collected over 19 sessions from six participants, totaling 23.8 hours of video. Runs were partitioned by scanning session, with the final run of each session held out for validation (19 runs, 6.7 h) and the remaining 93 runs used for training (17.1 h). Each frame is normalized per RGB channel to range [−1, 1], and resized to 120 × 90 pixels, preserving the 4:3 aspect ratio of the source. No data augmentation was applied.

Training examples are clips of 8 frames sampled at every 8th frame of the 30 fps source video, producing a tensor of shape 3 × 8 × 90 × 120 (channels × time × height × width). A clip beginning at source frame *i* comprises frames *i*, *i*+8, …, *i*+56, spanning 1.87 s within a reserved window of 2.13 s, matching the temporal resolution of fMRI data.

*Model*

The encoder and decoder of the VAE are composed of stacked residual 3D convolutional layers. The encoder consists of four layers with output dimensions of 64, 128, 256, and 512. Each block applied two 3D convolutions, each followed by batch normalization, with an activation function between them, and added a projection shortcut (1 × 1 × 1 convolution and batch normalization, matched in stride) before a final ELU activation function. The first layer used unit stride and a temporal kernel size of 1, acting frame-wise; the remaining layers used 3 × 3 × 3 kernels with stride 2 in both the temporal and spatial dimensions, applied in the first convolution of each block, downsampling with a factor of 8 in each dimension. A global average pool over the remaining temporal and spatial axes reduced the final feature map to a 512-dimensional vector.

The latent space was formed by linear mapping from the pooled encoder output to a 512-dimensional mean vector μ and log-variance vector log *σ*^2^, parameterizing a diagonal Gaussian posterior. For training stability, these values were bound to (−5, 5) by a soft clamp function defined as 5 tanh(x / 5). A latent embedding sampled from this distribution was fed to the decoder for reconstruction.

The decoder consisted of a fully connected layer that projected the latent sample to a 256 × 2 × 23 × 30 feature volume, followed by two residual 3D convolutional layers with upsampling and a final output head. For each layer, the frames are upsampled by trilinear interpolation to 4 × 45 × 60 at the first layer and 8 × 90 × 120 at the second layer to restore the original resolution. The same architecture as the encoder layer is used except that the stride size is kept at 1. The decoder mirrored the encoder resolutions with one fewer layer, with the omitted layer being absorbed into the initial fully connected layer.

The complete architecture of the VAE is summarized in Supplementary Table 1.

*Objective*

The model was trained by maximizing the evidence lower bound on the marginal log-likelihood of a clip (Kingma & Welling, 2013),

$$\log p\left( x \right)\geq E_{q\left( z | x \right)}\left[ \log p\left( x | z \right) \right]-D_{KL}(q\left( z | x \right)\|p(z))$$

in which q(z|x) is the approximate posterior produced by the encoder, p(x|z) the likelihood realized by the decoder, and p(z) = N(0, I) a fixed prior over a d-dimensional latent space with d = 512. The approximate posterior was taken to be diagonal Gaussian, q(z|x) = N(z; μ, diag(σ^2^)), so the divergence has the closed form

$$D_{KL}(q\left( z | x \right)\left\| p\left( z \right) \right)= -\frac{1}{2}\sum_{j=1}^{d} (1+log\sigma_{j}^{2}-\mu_{j}^{2}-\sigma_{j}^{2})$$

and the expectation was approximated with a single reparameterized sample per clip. Taking p(x|z) to be isotropic Gaussian with fixed variance reduces the expected log-likelihood to a squared error up to a constant. The KL divergence is weighted by a coefficient β. The final loss is

$$L=\left\| \hat{x}-x \right\|^{2}+\beta D_{KL}(q\left( z | x \right)\left\| p\left( z \right) \right), \beta=0.005$$

*Model Training*

Optimization used Adam at a constant learning rate of 1 × 10^-3^ (betas 0.9 and 0.999, eps 1 × 10^-8^, no weight decay), with gradients clipped to a global norm of 0.5. The batch size was 192, distributed across 3 NVIDIA RTX A6000 GPUs. One epoch covered all 925,778 training clips in 4,822 optimizer steps, and training proceeded for 242 epochs (1,164,102 optimizer steps, 461 h of wall-clock time). Models were implemented with PyTorch 1.13.1 (CUDA 11.6), Python 3.7.15, and PyTorch Lightning 1.8.5.

Voxelwise encoding models for VAE features

*Feature extraction*

The posterior mean of the latent embedding output of the VAE encoder was used as features for fitting voxelwise encoding models. For each run, the features were first extracted at 15 Hz by feeding 2.13 s clips of the stimulus to the encoder. The latent features were then resampled to 0.5 Hz using a 3-lobed Lanczos filter to match the 2 s TR of BOLD signals.

A finite impulse response (FIR) filter was used to capture effects of the hemodynamic response. To do so, the feature timeseries were delayed by 1, 2, 3, and 4 TRs, and were then concatenated to form 512 × 4 = 2048 input feature dimensions. As the BOLD signals were demeaned and the encoding models were fit without intercepts, the features were demeaned by the mean feature values from the training runs.

*Ridge regression*

For each voxel, the regularization parameter was selected by cross-validation over 50 logarithmically spaced values between 10^2^ and 10^15^. The cross-validation was performed on the training runs by holding out each single run as a validation fold. The model weights were estimated using himalaya.kernel_ridge.KernelRidgeCV (version=0.4.2) using the solver parameters of n_targets_batch=500, n_alphas_batch=5, and n_targets_batch_refit=100. Other parameters were set to their defaults.

*Weight normalization and aggregation*

For each voxel, the weights for each feature across the four FIR delays were first averaged. Since the norm of the weights were largely determined by the scale of regularization parameters rather than their effect size, and the regularization parameters differed across voxels, these raw weights were not directly comparable across voxels. Thus, the weight vectors were first normalized to unit norm and then rescaled by the normalized prediction performance of the encoding model on held-out test runs, quantified by the coefficient of determination (R^2^) between predicted and actual brain activity. The 1^st^ and 99^th^ percentiles of the prediction performance across voxels were first normalized to [0, 1] within participants, and then square-rooted. The model weights in each voxel were then rescaled by its normalized prediction performance value. This normalization placed model weights from each participant on the same scale to account for different functional SNR between participants. These rescaled weights were then projected to the fsaverage6 template and averaged across participants.

*Reconstructing preferred and anti-preferred visual experiences*

To reconstruct the preferred visual experiences for each functional region, the normalized group-average weights were first averaged across the functional region. Because magnitudes of the encoding model weights were out of distribution compared to the VAE embedding features (see Supplementary Figures 3 and 4), they could not directly be used to reconstruct visual experiences, as the decoder is sensitive to the magnitude of the embedding vector (see Supplementary Figures 5 and 6 for reconstructed visual experiences without rescaling). To make the encoding model weights usable for the decoder, they were rescaled to have a norm of 10, the minimal amount of rescaling needed to bring the norm of the weights into the native distribution for the VAE embedding vectors. Because the features were demeaned before fitting encoding models, the mean of the VAE features was added to this rescaled weight vector. Finally, this weight vector was given to the decoder to reconstruct the preferred visual experience. The same procedure was followed to reconstruct the anti-preferred visual experience with the negated average weight vector.
